# Annexin A2 Regulates Surfactant Dysfunction During Injurious Ventilation

**DOI:** 10.64898/2026.02.24.707549

**Authors:** Ian D. Bentley, Jonathan Fritz, Anusha Kapoor, R Duncan Hite, Samir N Ghadiali, Joshua A Englert

## Abstract

The Acute Respiratory Distress Syndrome (ARDS) is a life-threatening cause of respiratory failure, and patients who develop ARDS frequently require mechanical ventilation, which puts them at risk of developing ventilator induced lung injury (VILI). Both VILI and ARDS can induce pulmonary surfactant dysfunction, but the mechanisms are not known. Here we report a novel role for a phospholipid binding protein, Annexin A2 (AnxA2), in the regulation of surfactant composition and function following injurious ventilation. Wild type and AnxA2-/-mice were subjected to injurious ventilation and we found that AnxA2^-/-^ mice developed stiffer lungs following VILI that was not due to differences in barrier permeability or inflammation. Furthermore, we found that pulmonary surfactant from AnxA2^-/-^ mice had reduced surface tension lowering properties and that this was due to a reduction in 1-palmitoyl-2-oleoylphosphatidylglycerol, or POPG. Quantitative analysis of surface tension-surface area hysteresis loops obtained from surfactant isolated from AnxA2^-/-^ mice showed a defect in phase transitions during compression. In summary, Annexin A2 regulates surfactant function during injurious ventilation and may serve as a novel therapeutic target to prevent surfactant dysfunction in patients with ARDS who require mechanical ventilation.

## Introduction

The Acute Respiratory Distress Syndrome (ARDS) is a life-threatening cause of respiratory failure which affects approximately 200,000 people annually, and the most severe form of ARDS has a mortality rate approaching 50%.^1,2^ ARDS is characterized by excessive inflammation that develops following an initial insult such as sepsis, trauma, and pneumonia.^1^ This excess inflammation leads to breakdown of the alveolar-capillary barrier and flooding of the alveolar space with a protein and cell-rich edema fluid which impairs oxygenation.^1^ Patients with ARDS frequently require mechanical ventilation (MV), but during MV, lung cells are subjected to non-physiologic forces (e.g. atelectrauma, volutrauma) that cause ventilator-induced lung injury (VILI).^3^ Unfortunately, there are no molecularly targeted therapies to prevent or treat VILI in patients with ARDS.

One of the physiologic consequences of lung injury during ARDS and VILI is a reduction in lung compliance (i.e. increased lung stiffness).^4,5^ The decrease in compliance is due to a number of factors including noncardiogenic pulmonary edema and impaired activity of pulmonary surfactant.^1^ Pulmonary surfactant is a mixture of phospholipids and proteins secreted by type 2 alveolar epithelial (AT2) cells that lowers surface tension at the air-liquid interface to prevent alveolar collapse during respiration and promote gas exchange.^6^ Surfactant isolated from patients with ARDS and those requiring MV has been shown to have diminished surface tension lowering properties.^7,8^ Surfactant dysfunction can result from reduced synthesis or secretion of phospholipids and/or surfactant proteins, or increased surfactant degradation within the alveolar space.^6^ Although prior trials of exogenous surfactant replacement in ARDS have not demonstrated benefit,^9^ these studies were confounded by inadequate dosing and surfactant formulations that did not recapitulate the composition of endogenous surfactant.^10^ Drugs that augment the release of endogenous surfactant or prevent surfactant dysfunction have never been studied in patients.

Surfactant lipids are synthesized in AT2 cells and packaged into specialized organelles called lamellar bodies (LB), which traffic to the plasma membrane for release into the alveolar space.^6,11^ This pathway involves a calcium-sensitive, phospholipid-binding protein known as Annexin A2 (AnxA2).^12^ AnxA2 was originally identified in vascular endothelial cells, where it functions to promote fibrinolysis.^13,14^ AnxA2 is abundantly expressed in AT2 cells and has also been shown to play a role in secretion of vesicles from other secretory cell types, most notably neuroendocrine cells.^15^ In AT2 cells, AnxA2 has been shown to co-localize with LBs at the plasma membrane and has been implicated in LB fusion, but it’s exact role in LB release is not known.^12^ We hypothesized that AnxA2 plays a role in regulating surfactant function during VILI and may represent a novel therapeutic target for patients with ARDS who require MV. We found that AnxA2^-/-^ mice had decreased lung compliance during VILI. This difference was not due to changes in inflammation or barrier permeability; instead, it was associated with alterations in surfactant composition and function compared to wild-type mice.

## Methods

### Mouse Strains

Wild type C57BL/6 mice were obtained from Jackson laboratory (strain # 000664). AnxA2 global knockout (AnxA2^-/-^) mice were generously provided by the laboratory of Dr. Katherine Hajjar.^14^ Genotype was confirmed by qPCR using tail biopsies (Transnetyx) and western blotting using lung tissue. All mice were housed in pathogen free cages in the vivarium in The Davis Heart and Lung Research Institute at The Ohio State University (OSU). Experimental mice were between 8 and 14 weeks of age and provided with food and water ad lib. All animal experiments were performed under institutional review and approval of the protocol of the OSU Institutional Animal Care and Use Committee (protocol number: 2020A00000037).

### Murine Models of VILI

Mice were anesthetized with ketamine (100 mg/kg) and xylazine (10 mg/kg) and after induction of anesthesia, a tracheostomy cannula was inserted, and mice were placed on the Flexivent rodent ventilator (SCIREQ). Mice were subjected to four hours of injurious ventilation using a high tidal volume (12 cc/kg) and no positive end-expiratory pressure (PEEP) to induce two common types of VILI – volutrauma and atelectrauma. Throughout the ventilation procedure, mice were monitored using pulse oximetry (Starr Life Sciences) and warmed to 37ºC using the Harvard Apparatus Homeothermic System. Lung function measurements (e.g. compliance) were measured at baseline and every hour using the Flexivent system.^16^ Recruitment maneuvers were performed prior to lung function measurements to normalize lung volume. Sedation was maintained with alternating doses of ketamine and ketamine/xylazine as needed. After 4 hours of ventilation, mice were sacrificed with an anesthetic overdose. A bronchoalveolar lavage (BAL) was performed by instilling 1 cc of sterile 0.9% normal saline three times. BAL fluid was then centrifuged for 10 minutes at 400 g to pellet alveolar cells. Pelleted cells were treated with RBC lysis buffer (Thermo Fisher), counted using an automated cell counter (Bio-Rad), and cytospins were performed, followed by Hema-3 staining (Thermo Fisher) for manual differential cell counting. Spontaneously breathing control mice were sacrificed following induction of anesthesia but were not subjected to mechanical ventilation.

### Quantification of BAL Protein

Cell-free BAL fluid was subjected to bicinchoninic acid (BCA) assay (Thermo Fisher) to quantify protein in the BAL according to the manufacturer protocol.

### ELISA Protocol

Enzyme linked immunosorbent assays (ELISA) for mouse interleukin-6 (IL-6) and keratinocyte chemoattractant (KC) was performed with cell-free BAL fluid using mouse-specific ELISA detection and capture antibodies according to the manufacturer’s instructions (R+D Systems).

### Surfactant Isolation and Function Measurements

Cell-free BAL was ultracentrifuged at 40,000 g to isolate the small aggregate fraction (supernatant) and the large-aggregate fraction (pellet). The large aggregate (LA) was resuspended in a small volume of 0.9% saline. Phospholipids were extracted from the processed large and small aggregate fractions of pulmonary surfactant using a chloroform-methanol extraction.^17^ Total phosphorus content was then determined using a colorimetric assay following acid digestion of the phospholipids.^18^ Surface tension was measured in LA fractions following equalization of phospholipid content, using a custom-built constrained-drop surfactometer (CDS) as described previously.^19^

### Histology

Lungs were perfused with 10 mL of sterile phosphate buffered saline (PBS), and the left lung was inflated with 10% formalin to a pressure of 30 cmH2O prior to overnight fixation. Following fixation, lung tissue was stored in 70% ethanol and sent to Histowiz (Long Island City, NY) for embedding, sectioning, H&E staining and image preparation.

### Immunoblotting

Protein content of the LA fraction of murine surfactant was determined by BCA assay as detailed above. Samples were equalized in RIPA buffer (VWR) supplemented with protease and phosphatase inhibitors (Sigma-Aldrich) prior to boiling in Bolt LDS Buffer (Fisher Scientific) supplemented with 2.5% beta-mercaptoethanol (Bio-Rad).^20^ SDS-PAGE was performed using Nu-Page 12% Bis-Tris gels (Invitrogen) and proteins were transferred to a 0.2 um PVDF membrane (Bio-Rad). Membranes were incubated with primary and secondary antibodies as described in Supplemental Table 1. Bands were detected by chemiluminescence using SignalFire ECL Reagent (Cell Signaling Technology) and imaged on a ChemiDoc XRS+ System (Bio-Rad). Band density was then quantified using ImageJ software.

### Surfactant Phospholipid Identification

Phospholipids were extracted from a portion of LA following VILI by the Lipidomic Core at Wayne State University in accordance with the published protocols of the LipidMaps consortium.^21^ After addition of internal standards, total lipid extracts were prepared by extraction with methyl-*tert*-butyl ether.^17,22^ For analysis of phosphatidylcholine (PC) and phosphatidylethanolamine (PE), lipids were reconstituted in methanol-water-formic acid-ammonium formate and for analysis of phosphatidic acids (PA), phosphatidylglycerol (PG), and phosphatidylserine (PS), lipids were reconstituted in methanol-water-ammonium bicarbonate. All lipid samples were subjected to HPLC on Targa C8 (Higgins Analytical) and Multiple Reaction Monitoring (MRM) was used for the detection and quantification of each lipid molecule. Data were analyzed by LipidView software (ABSCIEX) for the identification and quantitation of lipids.

Phospholipids were extracted from a portion of LA from spontaneously breathing control mice by the Mass Spectrometry Core at the University of North Carolina using methyl-tert-butyl ether and ethanol. Following extraction, lipid analysis was performed using a Q Extractive Plus mass spectrometer (Thermo Fisher) coupled to an Acquity H-Class Liquid Chromatography System (Waters). Samples were analyzed in positive/negative switching ionization mode with top 5 data dependent fragmentation. Raw data was analyzed by LipidSearch (ThermoFisher) and lipids were analyzed by MS2 fragmentation.

### Biophysical Analysis of Lipid Dynamics within Pulmonary Surfactant

Loop morphological parameters were obtained using a custom-written MATLAB code using compression-expansion cycle number twenty from the LA fraction of pulmonary surfactant. Loop area was calculated by first smoothing experimental data to reduce noise and then integrating a cubic spline interpolation of this smoothed experimental data. Infection point magnitude was calculated by fitting expansion and compression data with a 6th-order polynomial, computing the 2nd derivative (e.g. “the rate of change of the rate of change”) and recording the maximum absolute value of the 2nd derivative as the inflection point magnitude.

### Statistical analysis

Data was analyzed using GraphPad Prism. Data are reported using column plots with mean +/-standard error of the mean. Sample size was determined by performing a pilot experiment and using the sample mean and standard derivation to calculate sample size required to achieve a power of 0.08 and α=0.05. All data was tested for normality using the Shapiro-Wilk test and for the presence of statistical outliers using the ROUT method with a Q threshold of 1%.^23^ Statistical outliers were excluded from the final statistical testing. Statistical comparisons between two groups were conducted using a Student’s t-test for parametric data, and the Mann-Whitney test for nonparametric data. Comparisons between multiple groups were conducted using a 1- or 2-way ANOVA test depending on the number of experimental variables. A p-value of <0.05 was considered statistically significant. When multiple comparisons were tested between groups, the Benjamini, Kreiger and Yekutieli method was used with a false positive rate of 10%.

## Results

### AnxA2^-/-^ mice have decreased lung compliance following injurious mechanical ventilation

Wild type (WT) and AnxA2^-/-^ mice were subjected to our VILI model. Prior to injury, AnxA2^-/-^ mice had less compliant (i.e. stiffer) lungs compared to WT controls (Fig 1A). Following VILI, the difference in lung compliance between genotypes became more exaggerated (Fig 1B). To investigate if the difference in lung stiffness was secondary to changes in barrier permeability, the concentration of protein in the BAL was measured and there was no difference between WT and AnxA2^-/-^ mice (Fig 1C). To evaluate whether the change in lung compliance was due to differences in alveolar inflammation, differential cell counts were performed, and pro-inflammatory cytokines and chemokines were measured in the BAL fluid. There was no difference in alveolar inflammation between WT and AnxA2^-/-^ mice following injury (Fig 1D, 1E, 1F). Following injury, lungs from WT and AnxA2^-/-^ mice were fixed in formalin and H&E staining was performed. High power images revealed the presence of interstitial thickening and evidence of alveolar edema in both WT and AnxA2^-/-^ mice without qualitative differences in injury (Fig 1G, 1H). These data demonstrate that the impaired lung compliance in AnxA2^-/-^ mice following VILI is not due to differences in alveolar permeability or inflammation.

**Figure 1.**
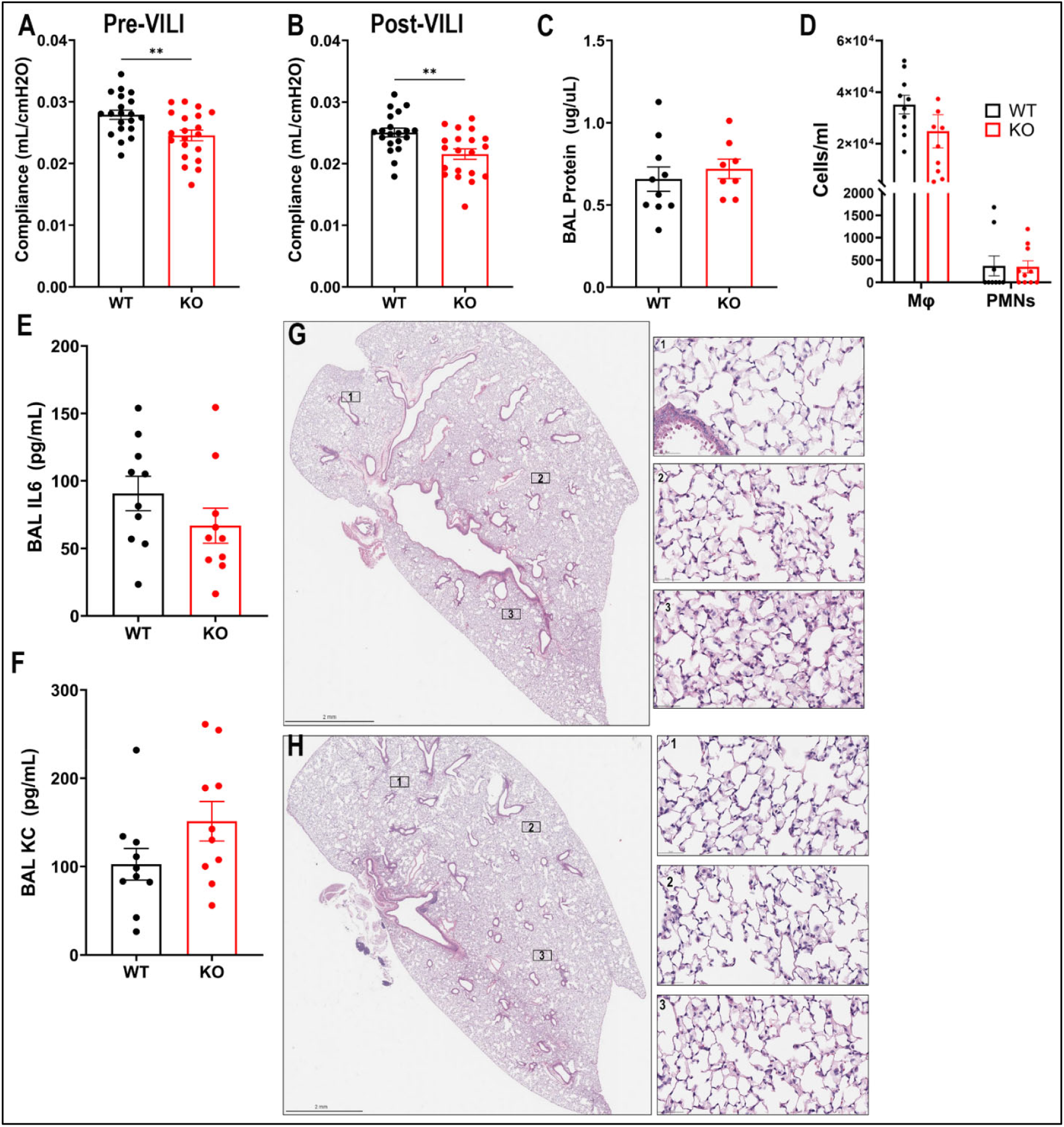
AnxA2^-/-^ mice are more susceptible to injurious mechanical ventilation. WT and AnxA2^-/-^ (KO) mice were subjected to 4 hours of injurious ventilation with 12 cc/kg tidal volume and 0 cmH2O PEEP to induce simultaneous volutrauma and atelectrauma. Lung compliance was measured before (A) and after (B) injurious ventilation. Following VILI, BAL was performed for measurement of barrier integrity (C) and differential cell counting (D). IL-6 (E) and KC (F) were measured in BAL fluid by ELISA. H&E staining was performed using formalin fixed lung tissue and representative images from a WT (G) and AnxA2^-/-^ (H) mice are shown following VILI. Numbered boxes in both G and H correspond to high-resolution (40x) panels to the right. ** p<0.01

### AnxA2^-/-^ mice have dysfunctional pulmonary surfactant following VILI

After determining that the increased susceptibility of AnxA2^-/-^ mice to VILI was not due to differences in alveolar barrier permeability or inflammation, we tested the hypothesis that AnxA2^-/-^ mice had impaired lung function following VILI due to impaired function of pulmonary surfactant. Surfactometry was performed and minimum surface tension was determined in the surface active large aggregate (LA) from WT and AnxA2^-/-^ mice.^19^ AnxA2^-/-^ mice had a significantly higher minimum surface tension (i.e. more dysfunctional surfactant) compared to WT mice. (Fig 2A). To investigate if impaired surfactant function in AnxA2^-/-^ mice was due to changes in phospholipid release or synthesis, the amount of surfactant phospholipid was measured. There were no significant differences in phospholipid content of the pulmonary surfactant or amounts in the LA or small aggregate fraction (Fig 2B, 2C, 2D). We then hypothesized that the differences in surfactant function following injury may be due to baseline differences in the lipid composition of surfactant. To test this hypothesis, we isolated surfactant from uninjured, spontaneously breathing WT and AnxA2^-/-^ mice, and found surfactant from AnxA2^-/-^ mice had an increased minimum surface tension (Fig 2G). VILI is known to impair surfactant function.^24,25^ Consistent with this, pulmonary surfactant isolated from spontaneous breathing WT and AnxA2^-/-^ mice exhibited lower minimum surface tension (i.e. more functional) than surfactant obtained after VILI. As observed following VILI, functional differences were not attributable to differences in phospholipid content (Fig 2H-J).

**Figure 2.**
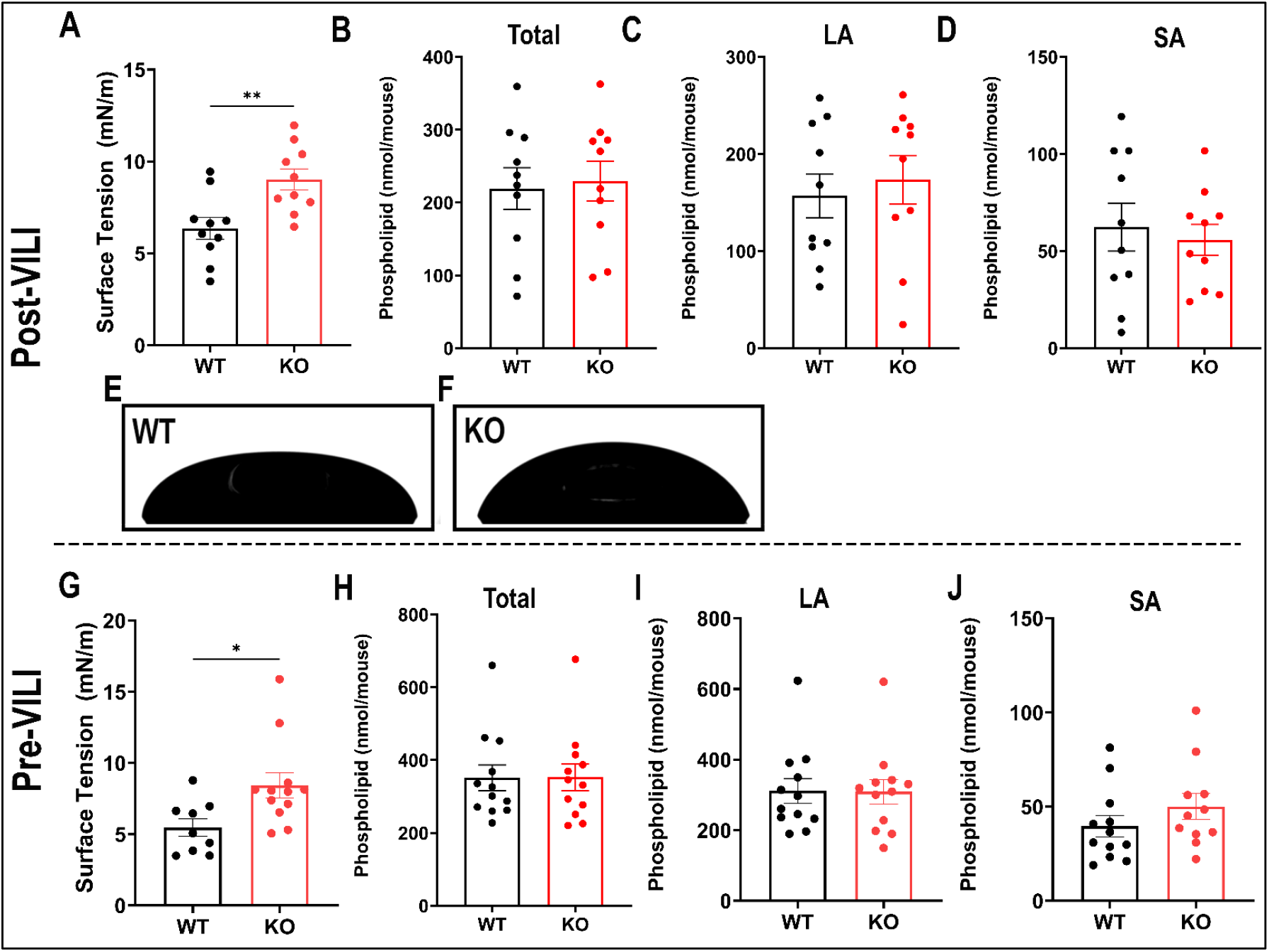
AnxA2^-/-^ mice have dysfunctional pulmonary surfactant following 4 hours of injurious ventilation. (A) Following VILI, the minimum surface tension of LA samples was measured using a constrained drop surfactometer (CDS). (B-D)Total phospholipid content was measured in whole pulmonary surfactant, the LA, and SA of pulmonary surfactant. (E,F) Representative images obtained from the CDS at minimum surface tension from a wild-type (WT) and AnxA2^-/-^ (KO) mouse following VILI. Pulmonary surfactant was isolated from uninjured, control mice and minimum surface tension was measured using the CDS (G) and phospholipid content was measured in total surfactant, the LA, and SA (H-J). *p<0.05, ** p<0.01.

### Surfactant proteins are not altered in AnxA2^-/-^ mice

In addition to phospholipids, the surface tension lowering properties of pulmonary surfactants are also mediated by surfactant proteins B and C (SP-B and SP-C).^6^ Both are secreted in lamellar bodies with phospholipids and are found in the surface-active LA fraction of pulmonary surfactant. To determine whether altered surfactant protein levels could account for the impaired surface tension lowering properties of surfactant from AnxA2^-/-^ mice, we performed immunoblotting of the LA fraction for both SP-B and SP-C (Fig 3A). Although surfactant protein expression was variable, there were no differences between AnxA2^-/-^ and WT mice (Fig 3B, 3C). Interestingly, AnxA2^-/-^ mice have less SP-C in the LA fraction prior to injury (Fig S1F), but this difference was not observed following injury. These data suggest that the impaired surfactant function in AnxA2^-/-^ mice is not mediated by changes in surfactant protein levels.

**Figure 3.**
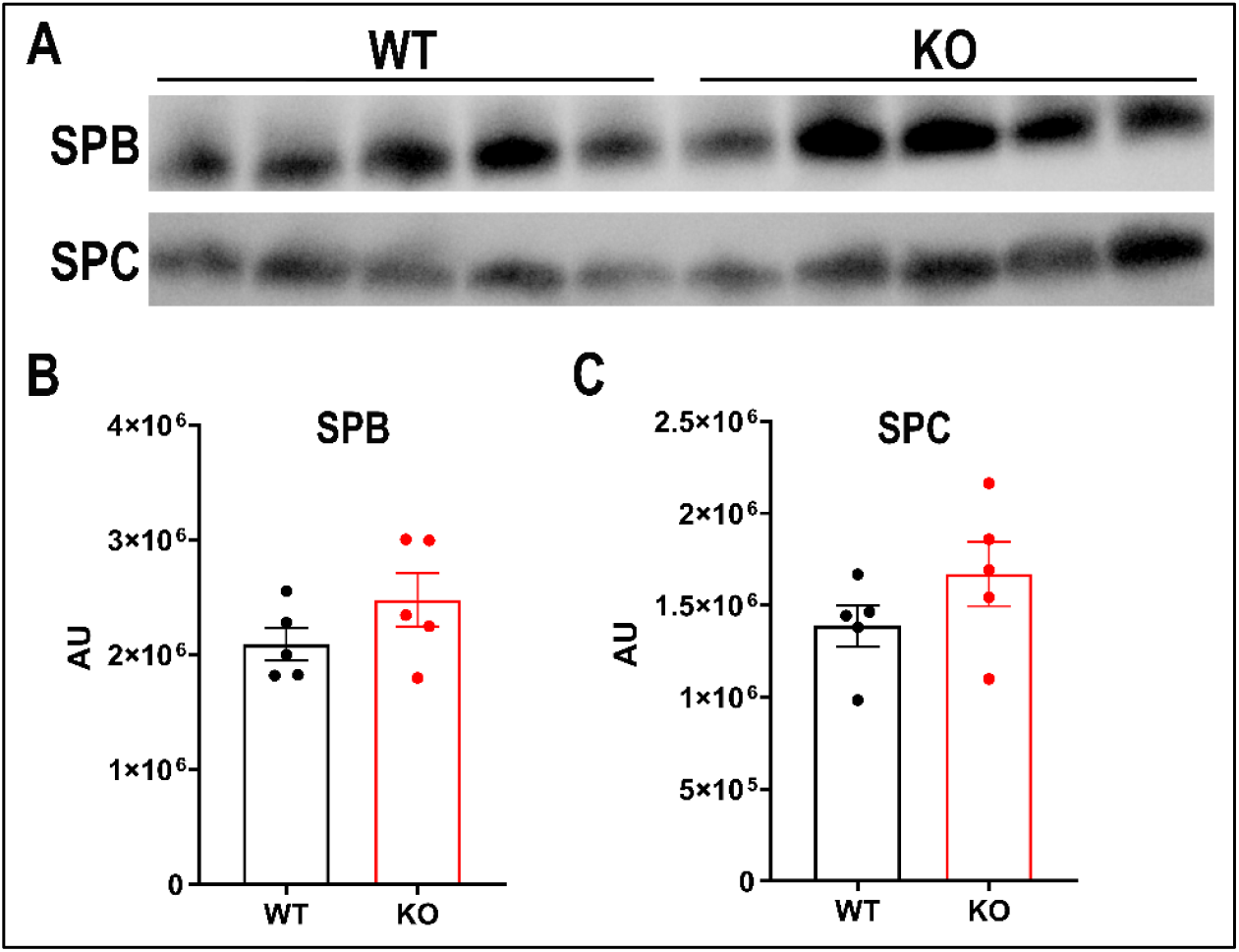
AnxA2^-/-^ mice surfactant protein composition is unchanged following 4 hour of injurious ventilation. Following VILI, LA was isolated from AnxA2^-/-^ and WT mice, and LA protein content was determined by BCA assay. Samples were equalized and subjected to SDS-PAGE followed by immunoblotting with primary antibodies (A). Band density is presented in arbitrary units (B, C).

### Lipidomic profiling reveals differences in AnxA2^-/-^ large aggregate phospholipid composition following VILI

Given that there were no differences in the total phospholipids or surfactant protein content in the LA fraction from WT and AnxA2^-/-^ mice, we assessed whether altered phospholipid composition could account for the differences in minimum surface tension between groups. Following VILI, lipids were extracted from the LA fraction for mass spectrometry analysis. Although there was a reduction in the amount of each individual phosphatidylcholine species in LA isolated from AnxA2^-/-^ mice, none of these reductions met significance after correction for multiple comparisons (Fig 4A). However, we did identify a significant reduction in three phosphatidylglycerol (PG) species (Fig 4B) after correction for multiple comparisons. We found that AnxA2^-/-^ mice have less PG (34:2), PG (34:1) and PG (32:1) in the LA fraction of pulmonary surfactant after VILI compared WT mice. Notably, one of these PG species, PG (34:1) (aka 1-palmitoyl-2-oleoylphosphatidylglycerol or POPG) is the predominant, non-phosphatidylcholine species in pulmonary surfactant.^6^ There were no significant differences identified in the other phospholipid classes that were measured: phosphatidylethanolamine, phosphatidylserine and phosphatidylinositol (Supplemental Fig 2). To confirm that the decrease in PG levels was not secondary to changes in phospholipid degradation within the alveolar space, phospholipase activity was measured in pulmonary surfactant isolated from AnxA2^-/-^ and WT mice following VILI. There were no differences in PG hydrolysis between AnxA2^-/-^ and WT mice (Fig 4C). In summary, these data suggest that changes in key PG species, including POPG, following VILI may mediate the change in lung compliance in AnxA2^-/-^ mice.

**Figure 4.**
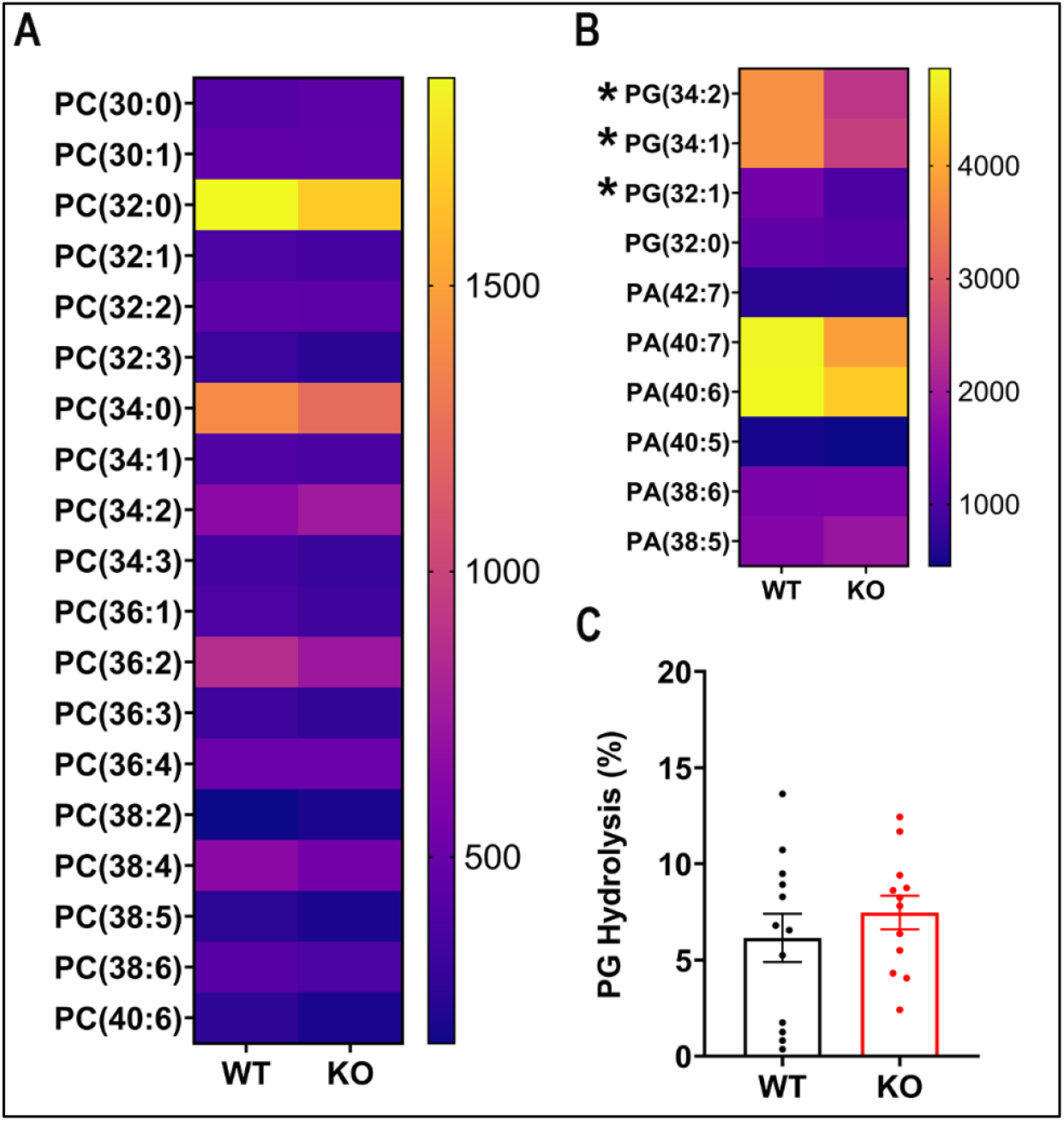
Following VILI, there is a reduction in three phosphatidylglycerol species in the LA of AnxA2^-/-^ mice. Following VILI, LA was isolated, and individual phospholipid species were identified by mass spectrometry. (A) Heat map of individual phosphatidylcholine species identified in LA following VILI. (B) Heat map of individual phosphatidylglycerol and phosphatidic acid species identified in LA following VILI. Units for the heat map are ng/mouse and data are expressed as a mean value over 5 replicates. Discoveries after correction for multiple comparisons are denoted with a *. (C) Following VILI, phospholipase A2 activity was measured in the small aggregate fraction of pulmonary surfactant and is reported as percent hydrolysis of phosphatidylglycerol.

### Genetic loss of AnxA2 induces surfactant dysfunction via compression defects at the air-liquid interface following injurious ventilation

To test the hypothesis that changes in POPG content in AnxA2^-/-^ pulmonary surfactant could explain the observed functional differences, we quantitatively analyzed changes in surface tension-surface area hysteresis loop morphology between genotypes. Surface tension-surface area loops from twenty successive compression-expansion cycles were generated using the constrained drop surfactometer (Fig 5A).^26^ While maximum surface tension did not differ between genotypes, LA from AnxA2^-/-^ mice demonstrated a higher minimum surface tension and a lower loop area (Fig 5B-D). During normal respiration, pulmonary surfactant must lower surface tension across a rapidly changing surface area as the alveolus inflates and collapses during each breath. To further quantify differences in loop morphology, we analyzed curvature during both are compression and expansion; where a second-order derivative was used to measure inflection magnitude. While there was no difference in inflection magnitude between WT and AnxA2^-/-^ samples during expansion, loops from AnxA2^-/-^ mice exhibited significantly reduced inflection magnitude during compression (Fig 5E-F). These results indicate that AnxA2 regulates surfactant biophysical properties during VILI by altering compression dynamics.

**Figure 5.**
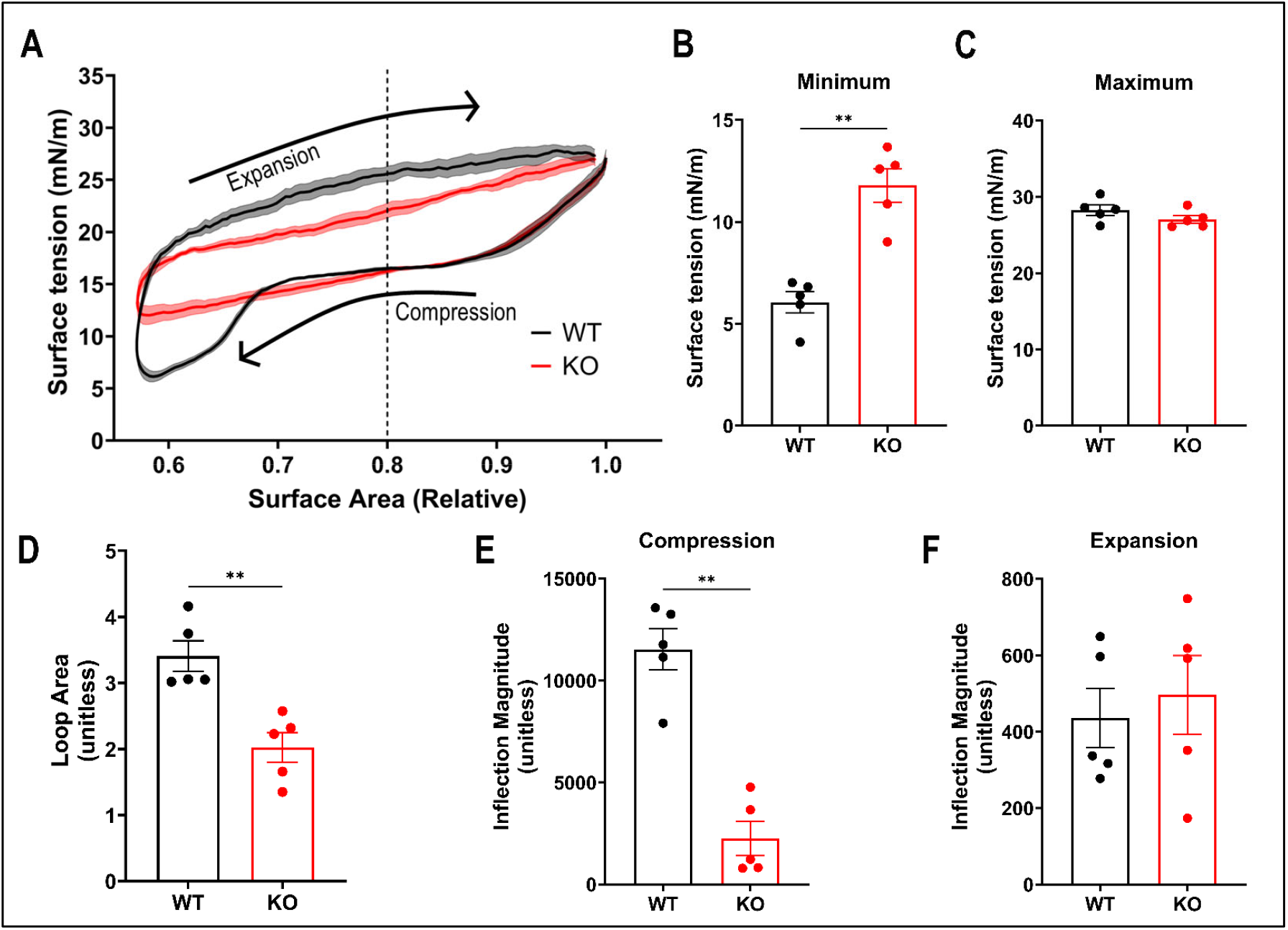
Following VILI, pulmonary surfactant from AnxA2^-/-^ mice demonstrates abnormal compression. (A) Surfactant hysteresis loops following 20 compression-expansion cycles on a CDS from LA fractions of WT or AnxA2^-/-^ after four hours of injurious ventilation (n = 5). Droplet surface area is normalized by indexing to the largest droplet surface area prior to compression. Data is presented as average surface tension (dark line) with standard error of the mean (shaded area) at any given relative surface area compression. (B-F) Loop morphological parameters obtained from Fig 5A: (B) Minimum surface tension. (C) Maximum surface tension. (D) Surface tension-surface area loop hysteresis area. The inflection magnitudes are calculated by finding the maximum absolute value of loop curvature (e.g. the second derivative) on compression (E) and expansion (F) below 80% compression (dashed line in (A)). **P < 0.01.

## Discussion

Patients with ARDS frequently require mechanical ventilation for life-support while clinicians treat the underlying cause of their lung injury. Positive pressure ventilation is known to cause or exacerbate underlying lung injury,^3^ and to date, prevention of VILI is limited to using “lung protective” tidal volumes and airway pressures.^27,28^ There are no pharmacologic interventions to treat or prevent VILI, partly because the molecular mechanisms by which non-physiologic forces drive injury are poorly understood. In this work, we report a novel role for AnxA2 in the regulation of surfactant function in a pre-clinical model of VILI. AnxA2^-/-^ mice were exposed to injurious ventilation and were found to have decreased respiratory system compliance compared to WT mice due to impaired surfactant function. We found that AnxA2 regulates levels of POPG in the alveolar space, and by utilizing a mathematical analysis of surface tension-area loops we found that AnxA2 regulates the surface tension lowering properties of pulmonary surfactant during compression.

Under homeostatic conditions, pulmonary surfactant lowers surface tension within the alveolar space and prevents alveolar collapse during tidal breathing.^6^ The absence of mature pulmonary surfactant at birth leads to the neonatal respiratory distress syndrome and surfactant function is also impaired in adults with pneumonia and ARDS.^8,29^ Pulmonary surfactant isolated from ARDS patients has been shown to have abnormal surface tension lowering properties which has been attributed to reduced phospholipid levels – including a reduction in phosphatidylglycerol.^30^ Interestingly, when our AnxA2^-/-^ mice were exposed to injurious ventilation, they developed a marked reduction in POPG, the most abundant phosphatidylglycerol species in surfactant.

Decreased levels of POPG in surfactant has been implicated in the development of neonatal respiratory distress syndrome,^31^ and increases in PG levels are associated with the development of mature pulmonary surfactant.^32^

Because the surface area of alveoli changes rapidly throughout the respiratory cycle, pulmonary surfactant must be able to rapidly adsorb and desorb to the air-liquid interface. During interfacial area compression, reorganization of surfactant phospholipid molecules at the air-liquid interface, and/or collapse of surfactant monolayers into multilayer structures result in lipid phase transitions.^33^ These phase transitions appear on compression-expansion isotherms as an inflection point and previous studies indicate that pulmonary replacement surfactants and bovine natural surfactants exhibit lipid phase transitions with large inflection points.^34^ Therefore, the lower inflection magnitude observed in the AnxA2^-/-^ loops during compression may be due to an inability to undergo phase transitions. Interestingly, POPG has been previously reported to facilitate the movement of surfactant phospholipids from the lipid reservoirs to the air-liquid interface during tidal respiration, a key element of these phase transitions.^35^

AnxA2 has previously been implicated in regulating lung compliance in uninjured mice,^36^ and consistent with this prior work, we observed that AnxA2^-/-^ mice have stiffer lungs in the uninjured state. This difference in compliance has been previously attributed to a defect in collagen 6 secretion from bronchial epithelial cells, which led to alterations in bronchiolar basement membranes and increased cell death.^36^ However, the physiologic mechanisms by which changes in the airway epithelium affect lung compliance remain unclear. Our study is the first to explore changes in surfactant function in these mice as an alternative mechanism of altered lung function. Altered surfactant surface tension lowering properties increase alveolar surface tension and increase lung stiffness.^37^ We observed changes in lung compliance and surfactant function in AnxA2^-/-^ mice both at baseline, and after ventilation, which suggests that changes in surfactant function may explain the observed differences in compliance.

Our study has several limitations. First, our murine VILI model induces a relatively modest injury, however we have intentionally chosen a tidal volume of 12 cc/kg to better reflect injurious tidal volumes that patients may be exposed to during routine clinical care.^2,38^ Second, while our quantitative analysis of biophysical properties of pulmonary surfactant following injury identified a defect in the ability of AnxA2^-/-^ pulmonary surfactant to undergo phase transitions during compression, this may not be the sole explanation for changes in minimum surface tension.

Detailed computational modeling of the complex fluid and surfactant transport dynamics that govern loop morphology will be needed to determine the precise biophysical mechanisms that increase minimum surface tension in AnxA2^-/-^ mice. Additional studies are also needed to determine the precise molecular mechanisms by which genetic deletion of AnxA2 decreases POPG levels. Here, we report a novel role for AnxA2 in the regulation of surfactant function during mechanical ventilation, by reducing the amount of POPG. Further investigation of this pathway may reveal therapeutic targets to promote endogenous surfactant function and limit injury in patients who require mechanical ventilation.

## Acknowledgements

We thank the University of North Carolina’s Department of Chemistry Mass Spectrometry Core Laboratory for their assistance with mass spectrometry analysis. Research reported in this publication was supported in part with funding by the University of North Carolina’s School of Medicine Office of Research. We thank the Wayne State University Lipidomic Core for their assistance with mass spectrometry analysis. Their core is supported by the Office of the Director of the National Institutes of Health under Award Numbers S10RR207926 and S10OD032292. Additionally, this publication was supported, in part, by the Ohio State University Clinical and Translational Science Institute and the National Center for Advancing Translational Sciences of the National Institutes of Health under Award Number UM1TR004548. The content is solely the responsibility of the authors and does not necessarily represent the official views of the National Institutes of Health. Finally, we thank the laboratory of Dr. Katheryn Hajjar at Weill Cornell Medical Center for generously providing Annexin A2 knockout mice, and the laboratory of Dr. Michael Beers at the University of Pennsylvania Perelman College of Medicine for generously providing Rabbit anti-mature surfactant protein C antibody.

## Author Contributions

IDB, JAE, SNG conceived and designed the project. IDB, JF, AK, RDH and SNG performed experiments and analyzed data. IDB wrote the manuscript. JAE provided overnight and coordination of the overarching research effort. All authors have reviewed and edited the manuscript.

## References

1. Thompson BT, Chambers RC, Liu KD. Acute Respiratory Distress Syndrome. Drazen JM, ed. N Engl J Med. 2017;377(6):562–572. doi:10.1056/NEJMra1608077

2. Bellani G, Laffey JG, Pham T, et al. Epidemiology, Patterns of Care, and Mortality for Patients With Acute Respiratory Distress Syndrome in Intensive Care Units in 50 Countries. JAMA. 2016;315(8):788–800. doi:10.1001/jama.2016.0291

3. Slutsky AS, Ranieri VM. Ventilator-Induced Lung Injury. N Engl J Med. 2013;369(22):2126–2136. doi:10.1056/NEJMra1208707

4. Seah AS, Grant KA, Aliyeva M, Allen GB, Bates JHT. Quantifying the Roles of Tidal Volume and PEEP in the Pathogenesis of Ventilator-Induced Lung Injury. Ann Biomed Eng. 2011;39(5):1505–1516. doi:10.1007/s10439-010-0237-6

5. Ashbaugh David G, Boyd Bigelow D, Petty Thomas L, Levine Bernard E. ACUTE RESPIRATORY DISTRESS IN ADULTS. The Lancet. 1967;290(7511):319–323. doi:10.1016/S0140-6736(67)90168-7

6. Agassandian M, Mallampalli RK. Surfactant phospholipid metabolism. Biochim Biophys Acta BBA - Mol Cell Biol Lipids. 2013;1831(3):612–625. doi:10.1016/j.bbalip.2012.09.010

7. Hallman M, Spragg R, Harrell JH, Moser KM, Gluck L. Evidence of lung surfactant abnormality in respiratory failure. Study of bronchoalveolar lavage phospholipids, surface activity, phospholipase activity, and plasma myoinositol. J Clin Invest. 1982;70(3):673–683.

8. Günther A, Siebert C, Schmidt R, et al. Surfactant alterations in severe pneumonia, acute respiratory distress syndrome, and cardiogenic lung edema. Am J Respir Crit Care Med. 1996;153(1):176–184. doi:10.1164/ajrccm.153.1.8542113

9. Günther A, Ruppert C, Schmidt R, et al. Surfactant alteration and replacement in acute respiratory distress syndrome. Respir Res. 2001;2(6):353–364. doi:10.1186/rr86

10. Lewis JF, Veldhuizen RAW. The Future of Surfactant Therapy during ALI/ARDS. Semin Respir Crit Care Med. 2006;27(4):377–388. doi:10.1055/s-2006-948291

11. Andreeva AV, Kutuzov MA, Voyno-Yasenetskaya TA. Regulation of surfactant secretion in alveolar type II cells. Am J Physiol-Lung Cell Mol Physiol. 2007;293(2):L259–L271. doi:10.1152/ajplung.00112.2007

12. Wang P, Chintagari NR, Gou D, Su L, Liu L. Physical and Functional Interactions of SNAP-23 with Annexin A2. Am J Respir Cell Mol Biol. 2007;37(4):467–476. doi:10.1165/rcmb.2006-0447OC

13. Flood EC, Hajjar KA. The annexin A2 system and vascular homeostasis. Vascul Pharmacol. 2011;54(3-6):59–67. doi:10.1016/j.vph.2011.03.003

14. Ling Q, Jacovina AT, Deora A, et al. Annexin II regulates fibrin homeostasis and neoangiogenesis in vivo. J Clin Invest. 2004;113(1):38–48. doi:10.1172/JCI19684

15. Gabel M, Delavoie F, Royer C, et al. Phosphorylation cycling of Annexin A2 Tyr23 is critical for calcium-regulated exocytosis in neuroendocrine cells. Biochim Biophys Acta BBA - Mol Cell Res. 2019;1866(7):1207–1217. doi:10.1016/j.bbamcr.2018.12.013

16. Bobba CM, Fei Q, Shukla V, et al. Nanoparticle delivery of microRNA-146a regulates mechanotransduction in lung macrophages and mitigates injury during mechanical ventilation. Nat Commun. 2021;12(1):1. doi:10.1038/s41467-020-20449-w

17. Bligh EG, Dyer WJ. A Rapid Method of Total Lipid Extraction and Purification. Can J Biochem Physiol. 1959;37(8):911–917.

18. Chen PS, Toribara TY, Warner Huber. Microdetermination of Phosphorus. Anal Chem. 1956;28(11):1756–1758. doi:10.1021/ac60119a033

19. Lee H, Fei Q, Streicher A, et al. mTORC1 is a mechanosensor that regulates surfactant function and lung compliance during ventilator-induced lung injury. JCI Insight. 2021;6(14):e137708. doi:10.1172/jci.insight.137708

20. Murthy A, Rodriguez LR, Dimopoulos T, et al. Activation of alveolar epithelial ER stress by β-coronavirus infection disrupts surfactant homeostasis in mice: implications for COVID-19 respiratory failure. Am J Physiol-Lung Cell Mol Physiol. 2024;327(2):L232–L249. doi:10.1152/ajplung.00324.2023

21. Quehenberger O, Armando AM, Brown AH, et al. Lipidomics reveals a remarkable diversity of lipids in human plasma. J Lipid Res. 2010;51(11):3299–3305. doi:10.1194/jlr.M009449

22. Matyash V, Liebisch G, Kurzchalia TV, Shevchenko A, Schwudke D. Lipid extraction by methyl-tert-butyl ether for high-throughput lipidomics. J Lipid Res. 2008;49(5):1137–1146. doi:10.1194/jlr.D700041-JLR200

23. Motulsky HJ, Brown RE. Detecting outliers when fitting data with nonlinear regression – a new method based on robust nonlinear regression and the false discovery rate. BMC Bioinformatics. 2006;7(1):123. doi:10.1186/1471-2105-7-123

24. Gregory TJ, Longmore WJ, Moxley MA, et al. Surfactant chemical composition and biophysical activity in acute respiratory distress syndrome. J Clin Invest. 1991;88(6):1976–1981. doi:10.1172/JCI115523

25. Maruscak AA, Vockeroth DW, Girardi B, et al. Alterations to surfactant precede physiological deterioration during high tidal volume ventilation. Am J Physiol-Lung Cell Mol Physiol. 2008;294(5):L974–L983. doi:10.1152/ajplung.00528.2007

26. Yu LMY, Lu JJ, Chan YW, et al. Constrained sessile drop as a new configuration to measure low surface tension in lung surfactant systems. J Appl Physiol. 2004;97(2):704–715. doi:10.1152/japplphysiol.00089.2003

27. Amato MBP, Meade MO, Slutsky AS, et al. Driving pressure and survival in the acute respiratory distress syndrome. N Engl J Med. 2015;372(8):747–755. doi:10.1056/NEJMsa1410639

28. Acute Respiratory Distress Syndrome Network, Brower RG, Matthay MA, et al. Ventilation with lower tidal volumes as compared with traditional tidal volumes for acute lung injury and the acute respiratory distress syndrome. N Engl J Med. 2000;342(18):1301–1308. doi:10.1056/NEJM200005043421801

29. Griese M, Birrer P, Demirsoy A. Pulmonary surfactant in cystic fibrosis. Eur Respir J. 1997;10(9):1983–1988. doi:10.1183/09031936.97.10091983

30. Pison U, Seeger W, Buchhorn R, et al. Surfactant Abnormalities in Patients with Respiratory Failure after Multiple Trauma. Am Rev Respir Dis. 1989;140(4):1033–1039. doi:10.1164/ajrccm/140.4.1033

31. Haumont D, Rossle C, Clercx A, et al. Modifications of Surfactant Phospholipid Pattern in Premature Infants Treated with Curosurf: Clinical and Dietary Correlations. Biology of the Neonate. 1992;61:37–43.

32. Hallman M, Gluck L. Phosphatidylglycerol in lung surfactant. III. Possible modifier of surfactant function. J Lipid Res. 1976;17(3):257–262. doi:10.1016/S0022-2275(20)36982-0

33. Possmayer F, Zuo YY, Veldhuizen RAW, Petersen NO. Pulmonary Surfactant: A Mighty Thin Film. Chem Rev. 2023;123(23):13209–13290. doi:10.1021/acs.chemrev.3c00146

34. Zhang H, Fan Q, Wang YE, Neal CR, Zuo YY. Comparative study of clinical pulmonary surfactants using atomic force microscopy. Biochim Biophys Acta BBA - Biomembr. 2011;1808(7):1832–1842. doi:10.1016/j.bbamem.2011.03.006

35. Liekkinen J, Enkavi G, Javanainen M, Olmeda B, Pérez-Gil J, Vattulainen I. Pulmonary Surfactant Lipid Reorganization Induced by the Adsorption of the Oligomeric Surfactant Protein B Complex. J Mol Biol. 2020;432(10):3251–3268. doi:10.1016/j.jmb.2020.02.028

36. Dassah M, Almeida D, Hahn R, Bonaldo P, Worgall S, Hajjar KA. Annexin A2 mediates secretion of collagen VI, pulmonary elasticity and apoptosis of bronchial epithelial cells. J Cell Sci. 2014;127(4):828–844. doi:10.1242/jcs.137802

37. Ingenito EP, Tsai LW, Majumdar A, Suki B. On the Role of Surface Tension in the Pathophysiology of Emphysema. Am J Respir Crit Care Med. 2005;171(4):300–304. doi:10.1164/rccm.200406-770PP

38. Needham DM, Colantuoni E, Mendez-Tellez PA, et al. Lung protective mechanical ventilation and two year survival in patients with acute lung injury: prospective cohort study. The BMJ. 2012;344:e2124. doi:10.1136/bmj.e2124

39. Beers MF, Bates SR, Fisher AB. Differential extraction for the rapid purification of bovine surfactant protein B. Am J Physiol-Lung Cell Mol Physiol. 1992;262(6):L773–L778. doi:10.1152/ajplung.1992.262.6.L773

