## Supplemental Table 1 for "Annexin A2 Regulates Surfactant Dysfunction During Injurious Ventilation"

| **Supplemental Table 1: Antibodies used for immunoblotting** | | | |
| --- | --- | --- | --- |
| **Antibody** | **Manufacturer** | **Catalog Number** | **Dilution Used** |
| Rabbit Anti-Mature SP-B | Generously provided by Michael Beers^39^ | n/a | 1:3000 in 5% Milk/TBST |
| Rabbit Anti-Mature SP-C | Seven Hills | WRAB-76694 | 1:3000 in 5% Milk/TBST |
| Goat anti-rabbit HRP-IgG | Cell Signaling Technology | 7074S | 1:5000 in 1% Milk/TBST |
