## Supplemental Figure 1 for "Annexin A2 Regulates Surfactant Dysfunction During Injurious Ventilation"

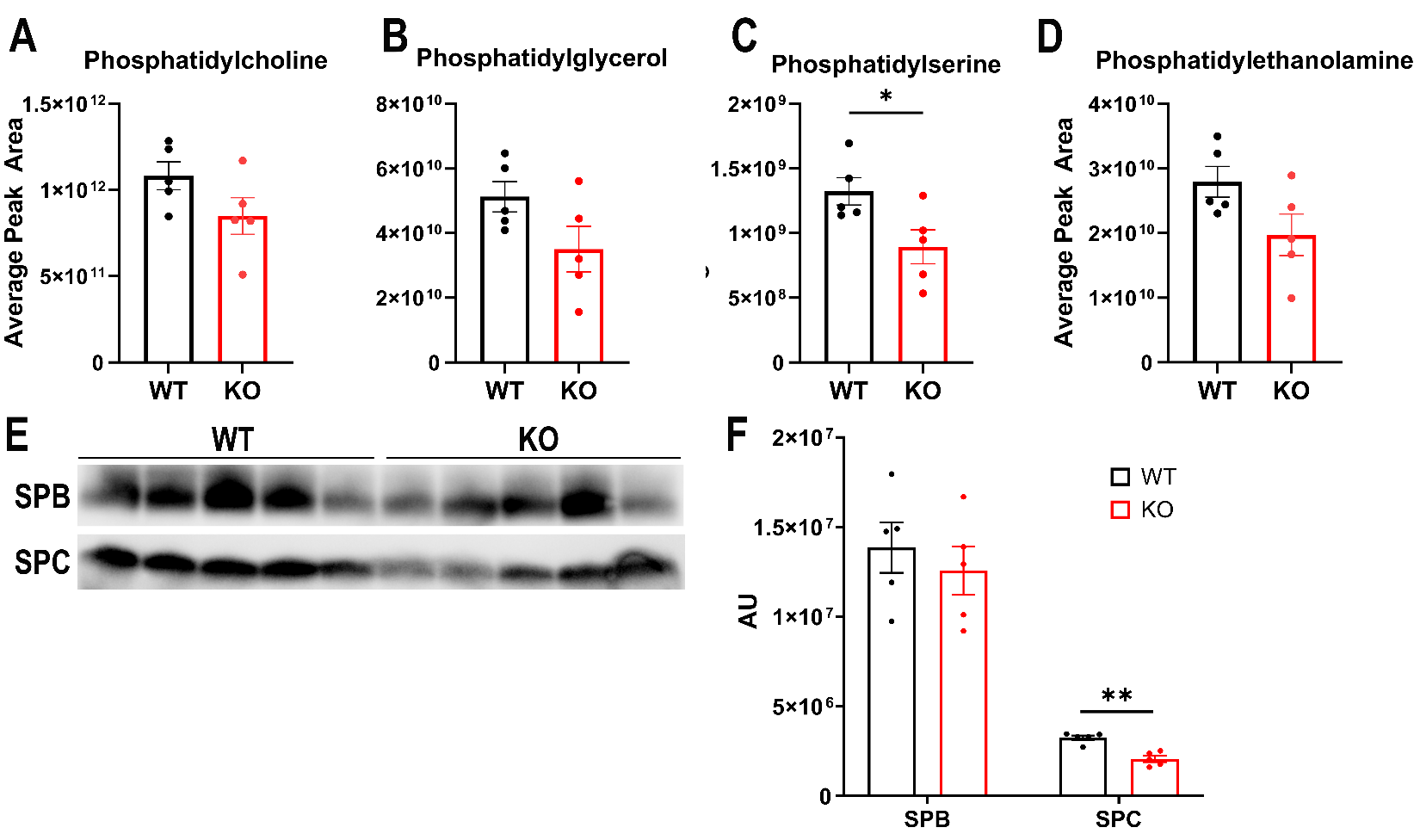

**Supplemental Figure 1:** **Compositional profiling of pulmonary surfactant isolated from spontaneously breathing WT and AnxA2^-/-^ mice.** Individual phospholipid classes within the LA were identified by mass spectrometry (A-D). Total protein content of the LA was measured by BCA assay, equalized and subjected to SDS-PAGE for immunoblotting by primary antibodies (E). Band density is presented in arbitrary units (F). * P<0.05, ** p<0.01
