## Supplemental Figure 2 for "Annexin A2 Regulates Surfactant Dysfunction During Injurious Ventilation"

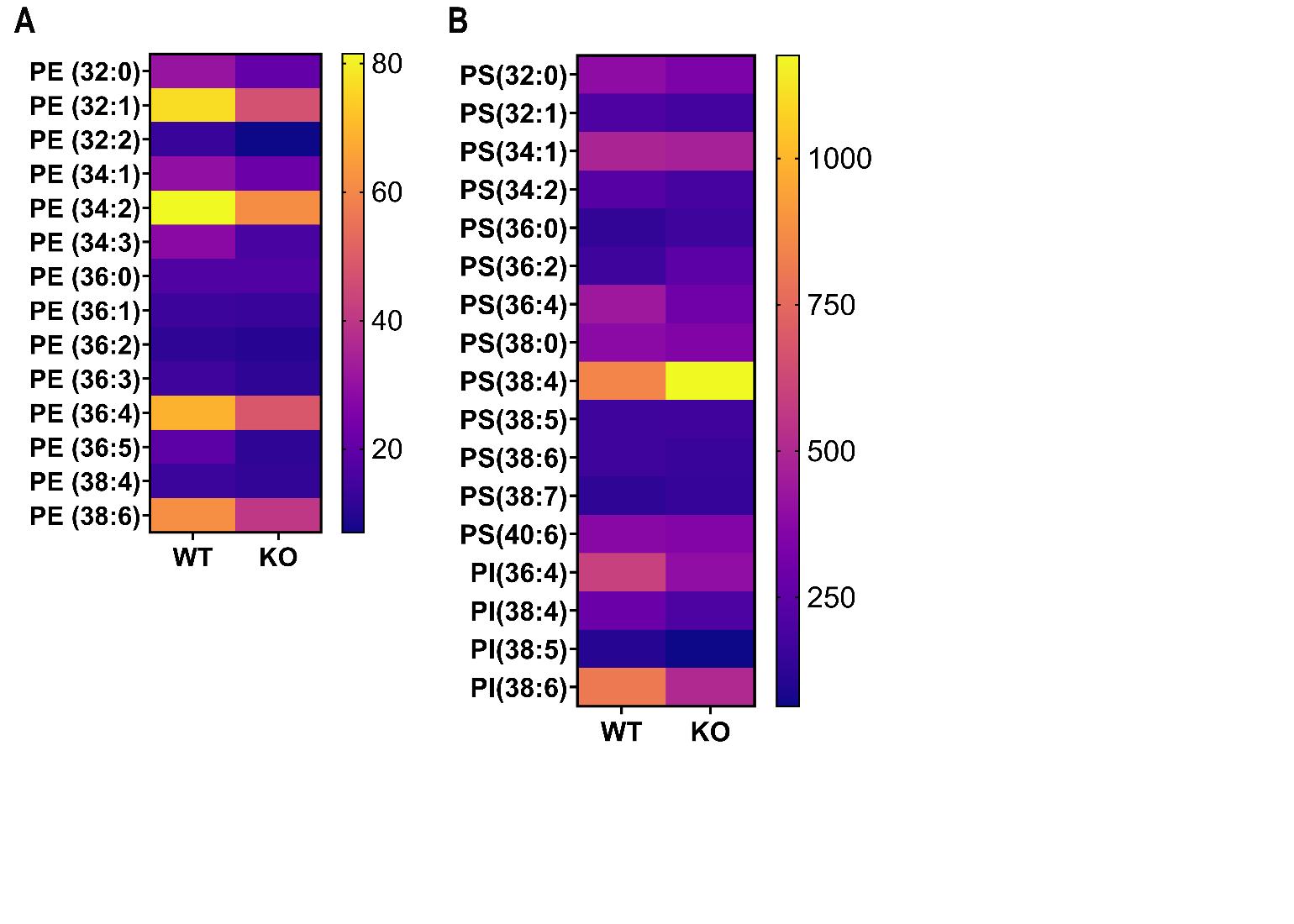

**Supplemental Figure 2.** **Identification of minor phospholipid species in the large aggregate of WT and AnxA2^-/-^ mice.** Following VILI, LA was isolated, and individual phospholipid species were identified by mass spectrometry. (A) Heat map of individual phosphatidylethanolamine species identified in LA following VILI. (B) Heat map of individual phosphatidylserine and phosphatidylinositol species identified in LA following VILI. Units for the heat map are ng/mouse and data are expressed as a mean value over 5 replicates. Significance testing was performed after removal of low-abundance phospholipid species by the Benjamini, Kreiger and Yekutieli method of correcting for multiple comparisons with an allowed false positive rate of 10%.
