## Supplemental Figure 3 for "Annexin A2 Regulates Surfactant Dysfunction During Injurious Ventilation"

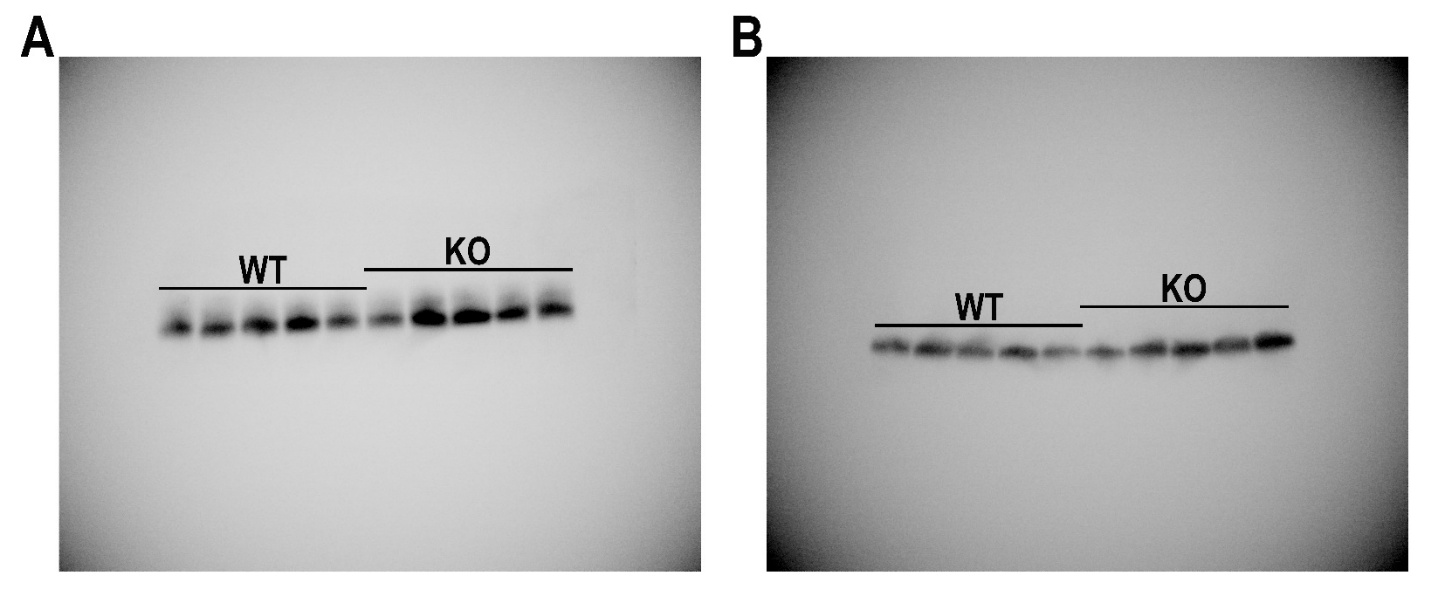

**Supplemental Figure 3. Uncropped western blots, presented in their cropped form in Figure 3.** Following VILI, LA was isolated from AnxA2^-/-^ and WT mice, and LA protein content was determined by BCA assay. Samples were then equalized and subjected to SDS-PAGE followed by immunoblotting with anti-mature surfactant protein B (A) and anti-mature surfactant protein C (B) antibodies.
